# Myelin Decompaction in Mice Given Anesthetics during Magnetic Resonance Imaging

**DOI:** 10.1101/2025.07.29.667499

**Authors:** Charlotte Best, Harikesh Dubey, Sheng Liu, Antonio White, Sravya Jayam, Christiane L. Mallett, Rebecca C. Knickmeyer

**Author notes:** Corresponding author: Rebecca C. Knickmeyer Corresponding author’s.

## Abstract

The objective of this secondary analysis of a prior investigation was to determine if prolonged exposure to the anesthetics isoflurane and dexmedetomidine during MRI was associated with a higher proportion of axons with myelin decompaction. 16 mice underwent an MRI protocol in which they had prolonged exposure to isoflurane and dexmedetomidine, while 10 mice did not undergo this protocol. All mice were sacrificed and electron microscope images were taken of various brain regions including the right prefrontal cortex (anterior cingulate and prelimbic area), the nucleus accumbens, the amygdala, and the ventral hippocampus.. Proportion of decompacted axons was calculated for each mouse, and an inter-rater reliability score of 80% was achieved. Welch’s t-tests were used to test the hypothesis that mice undergoing MRI with prolonged anesthesia had greater levels of myelin decompaction than mice that did not experience prolonged anesthesia. Mice with prolonged anesthetic exposure during MRI had significantly higher proportions of decompacted axons than mice that did not experience prolonged anesthesia (p-value of 0.00003642). Prolonged exposure to anesthetics, particularly isoflurane, may be associated with myelin decompaction. These findings, if replicated, have potential to impact future anesthesia use in clinical work and scientific research.

**Significance Statement:** Prolonged exposure to the anesthetics isoflurane and dexmedetomidine during brain imaging may lead to myelin decompaction in adult rodents. Myelin is a protective sheath around nerve fibers that ensures efficient transmission of electrical signals in the nervous system. Decompaction of myelin can disrupt these signals, potentially leading to neurological issues. This discovery is significant because it highlights potential risks associated with these anesthetics, which are commonly used in fMRI studies of rodents and in veterinary and medical procedures.

## Introduction

Myelin is essential for proper nervous system functioning in humans and animals (Morrell and Quarles, 1999). The myelin sheath, consisting of several layers of fatty membranes composed of cholesterol and glycolipid, wraps around the axon to insulate it, increasing the speed of conductance of action potentials along the axon fiber (Poitelon et al., 2020). If the myelin sheath is damaged, a variety of issues can result from improper signal conductance. More specifically, a type of damage called myelin decompaction occurs when individual sheaths become loose and expand outward away from the axon. This can lead to reduced motor coordination and functional connectivity in rodents (Asleh et al., 2020).

Myelin is especially susceptible to damage during development, which occurs from birth until around postnatal day 60 in mice (Hammelrath et al., 2016). During this critical period, oligodendrocytes proliferate throughout the brain to promote myelin formation and stability (Simons and Nave, 2015). If oligodendrocyte activity is disrupted, myelination may also be disrupted. This can happen in many ways, but one of particular interest is administration of anesthetics such as isoflurane. Isoflurane is a volatile anesthetic used for induction and maintenance of general anesthesia and is currently used in animals and humans (Hawkley et al., 2025). Previous studies have shown that isoflurane administered during development disrupts oligodendrocyte activity, leading to altered myelination lasting into adulthood (Kang et al., 2017). Isoflurane administered during development also result in neuroapoptosis, another mechanism that reduced neuronal functioning (Zhao et al., 2020). Interestingly, other anesthetics are thought to have a protective effect against myelination disruption. In particular, dexmedetomidine, a highly selective α2 adrenergic receptor agonist, promotes survival and maturation of oligodendrocyte progenitors and enhances myelination in a rodent model of neonatal hypoxic-ischemic brain injury (Xue et al., 2025). α2 adrenergic activity in the frontal cortex is positively correlated with oligodendrocyte density and mRNA levels of several oligodendrocyte markers including PLP1 and OLIG2. The same study found that the GO pathway for myelination was significantly overrepresented among genes correlated with α2 adrenergic activity in frontal cortex (Kim et al., 2012).

While there is a substantial amount of research on the impacts of isoflurane and dexmedetomidine on neuronal function and myelination during mouse development, there is less research on the effect of these anesthetics on myelin in adulthood. One study has reported motor decrements and altered white matter integrity in adult mice after prolonged isoflurane exposure, but to our knowledge, this is the only published study exploring this topic (Bajwa et al., 2019).

While conducting a study on the impact of microbiome communities on mouse behavior, brain imaging phenotypes, and axonal phenotypes, we unexpectedly discovered substantial myelin sheath decompaction in adult animals exposed to isoflurane and dexmedetomidine during magnetic resonance imaging (MRI). As isoflurane and dexmedetomidine are used for many experimental protocols involving mice, including studies of neurobiological and behavioral outcomes, we wish to draw attention to this phenomenon. Our findings may also have clinical implications as isoflurane and dexmedetomidine are widely used in both medical and veterinary practice, affecting both patients and health care workers who experience occupational exposures. Furthermore, isoflurane is being actively investigated as a treatment for depression, and our findings may have implications for those studies as well (Breault et al., 2025).

## Methods

### Overview

This is a secondary data analysis study to determine whether myelin decompaction differs between adult mice exposed to anesthetics during MRI and mice without prolonged exposure to anesthetics. The objective of the parent study was to determine if variation in human infant gut microbiomes influenced neurodevelopment using humanized mouse models (Dubey et al., 2023). Microbial communities derived from human infants were transplanted into pregnant germ-free mice. Offspring were then monitored and underwent several behavioral paradigms, and a subset underwent MRI with anesthetic exposure prior to humane euthanization at 9-10 weeks postnatal age. After euthanasia, myelin phenotypes for all animals were assessed using transmission electron microscopy. Transplantation and subsequent tests were conducted in 3 cohorts across a period of approximately 11 months (August 2020 through June 2021).

### Study Mice

Pregnant Swiss-Webster germ-free (GF) mice were generated by the University of Michigan Germ-Free (GF) Mouse Facility. All recipient mice were between 6 and 8 weeks of age at the start of pregnancy and were randomly selected for fecal slurry inoculations. Fecal slurries were administered by oral gavage on embryonic day 10 (ED10) between 1:00-2:00pm as a single dose of 200µL. Group 1 (HUM1 group) was inoculated with a mixed human fecal slurry derived from infants with relatively high levels of *Bifidobacterium*. Group 2 (HUM2 group) was inoculated with a mixed human fecal slurry derived from infants with relatively high levels of *Bacteroides*. Group 3 (SPF control) received fecal slurry from specific pathogen free mice (SPF group) and Group 4 (GF group) received autoclaved fecal slurry from SPF mice. For the first three groups (HUM1, HUM2, and SPF), pregnant mice were transferred to the Michigan State University Animal Care Facility between Day 14 and Day 20 of pregnancy in sterile containers. The GF group remained at University of Michigan through weaning and the offspring were transferred to MSU around 5 weeks of age.

Campus Animal Resources (CAR) provided veterinary care, daily husbandry, and health checks for all animals. Animals were housed in a special room dedicated to mice carrying human microbiotas. Each litter occupied a single cage, with their mother, until weaning. After weaning animals were group housed with littermates of the same sex (4-5 animals per cage) until they were about 8 weeks of age, when we conducted a sucrose preference test, which requires single housing. Animals remained singly housed until the end of the study. Over the course of 8 weeks, the mice underwent the following behavioral tests: elevated plus maze, light/dark preference, open field, three-chamber sociability test, novel object recognition test, sucrose preference test, carmine red test, and stress reactivity test. When animals were around 9-10 weeks of age, a subset underwent MRI imaging under prolonged anesthesia prior to humane euthanasia. The other animals were humanely euthanized at the same age, but without MRI imaging and prolonged exposure to anesthetics. These animals were briefly exposed to anesthesia as part of the humane euthanasia protocol.

### MRI and Anesthesia Protocol

Prior to the MRI, mice were placed in an induction chamber filled with 2-5% isoflurane with an oxygen flow rate of about 1 L/min. Once the mouse was deeply anesthetized, operationalized as no motion and slow, deep, even breaths, it was removed from the chamber. The eyes were covered with ophthalmic ointment to prevent them from drying. The mouse was moved to a nose cone lying on top of a warm water circulating pad (covered by a clean bench diaper or wypal).

The nose cone continued to deliver isoflurane at 0.5-2.5%. One cannula was inserted in the subcutaneous tissue of the left flank to deliver dexmedetomidine during the scan; a second cannula was inserted into the subcutaneous tissue of the right flank to deliver warm saline injections of 100-300 uL °every 30 minutes throughout the scan. Monitoring equipment was attached, consisting of a rectal temperature probe and a pneumatic pressure sensor placed on the chest. After fixing the equipment with adhesive tape, the animal was transferred to a MRI-compatible mouse bed and MRI was performed in a 7T MRI (Bruker, Billerica, MA).

Dexmedetomidine 0.5mg/ml stock solution was diluted with saline to generate a bolus dose of 0.05 mg/kg at 0.0 mg/mL which was delivered via subcutaneous injection. After 15 minutes, we began infusion of dexmedetomidine at 0.02 mg/mL in uL/min through the cannula inserted in the left flank subcutaneous tissue. Isoflurane was reduced gradually over the next 10-25 minutes. Rectal temperatures were kept around 36.5 ± 1 °C using a pad heated by circulating water. Isoflurane was adjusted to maintain respiration between 30 to 120 RR/min. fMRI studies require a RR/min around 90 to 120; while structural scans can be conducted at lower RR/min. Anesthesia was generally maintained for 2-3 hours. Figure 1 displays spaghetti plots of breathing rate, temperature, and isoflurane concentration during this time period for each mouse.

**Figure 1.**
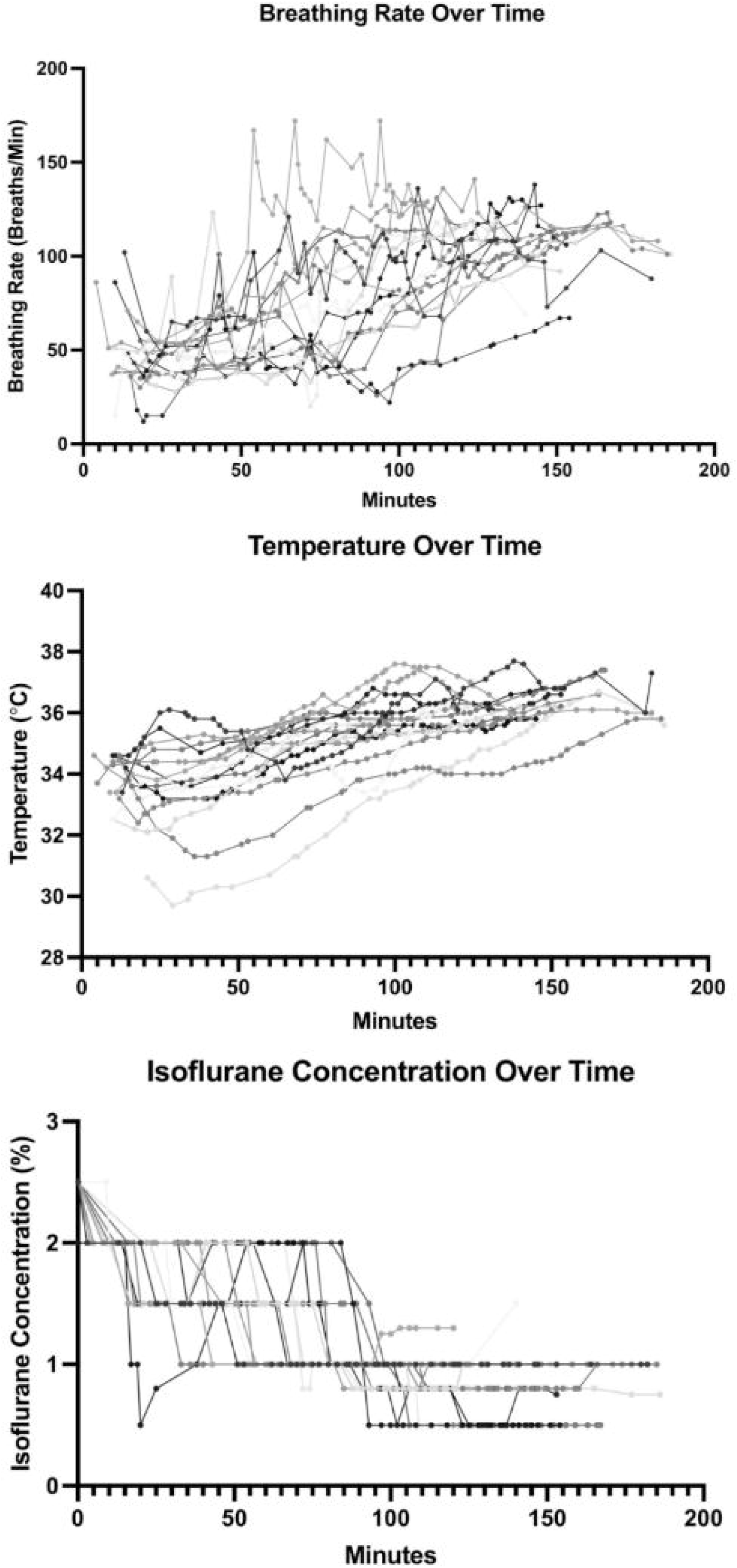
Spaghetti plots showing breathing rate, temperature, and isoflurane concentration over time in mice during MRI procedure. Each line indicates a separate mouse.

### Euthanasia and Transmission Electron Scanning

After MRI, mice immediately underwent cardiac perfusion and were euthanized via cervical dislocation. Mice that didn’t undergo MRI were sacrificed in the same manner. All mice were anesthetized using isoflurane prior to cardiac perfusion. This means that the non-MRI group was exposed to isoflurane, but for significantly less time than the mice in the MRI group.

After being euthanized, the brains were removed and a Zivic brain slicer tool, scalpel blades, and 1.5mm and 1.2mm tissue punches were used to collect samples of the right prefrontal cortex (anterior cingulate and prelimbic area), the nucleus accumbens, the amygdala, and the ventral hippocampus. Each region was stored in 4% paraformaldehyde for electron microscopy of axonal phenotypes.

After primary fixation, samples were washed with 0.1M phosphate buffer and postfixed with 1% osmium tetroxide in 0.1M phosphate buffer and further dehydrated in a gradient series of acetone. Samples were embedded in Spurr resin. For each specimen 70 nm thin sections were obtained from the polymerized block using a Power Tome Ultramicrotome (RMC, Boeckeler Instruments. Tucson, AZ). The thin sections were stained with uranyl acetate and lead citrate stain. Further, images were examined using a JEOL 1400 Flash Transmission Electron Microscope (Japan Electron Optics Laboratory, Japan) at an accelerating voltage of 100kV. Electron micrographs were obtained from the regions described above.

### EM Image Analysis

EM images were analyzed in ImageJ. Axons were labeled as “decompacted” (DC) or “non- decompacted” (N) by two raters (Figure 2). Raters were unaware of each other’s labels, and an inter-rater reliability rate of 81% was achieved. A total of 1,658 axons were labelled.

**Figure 2.**
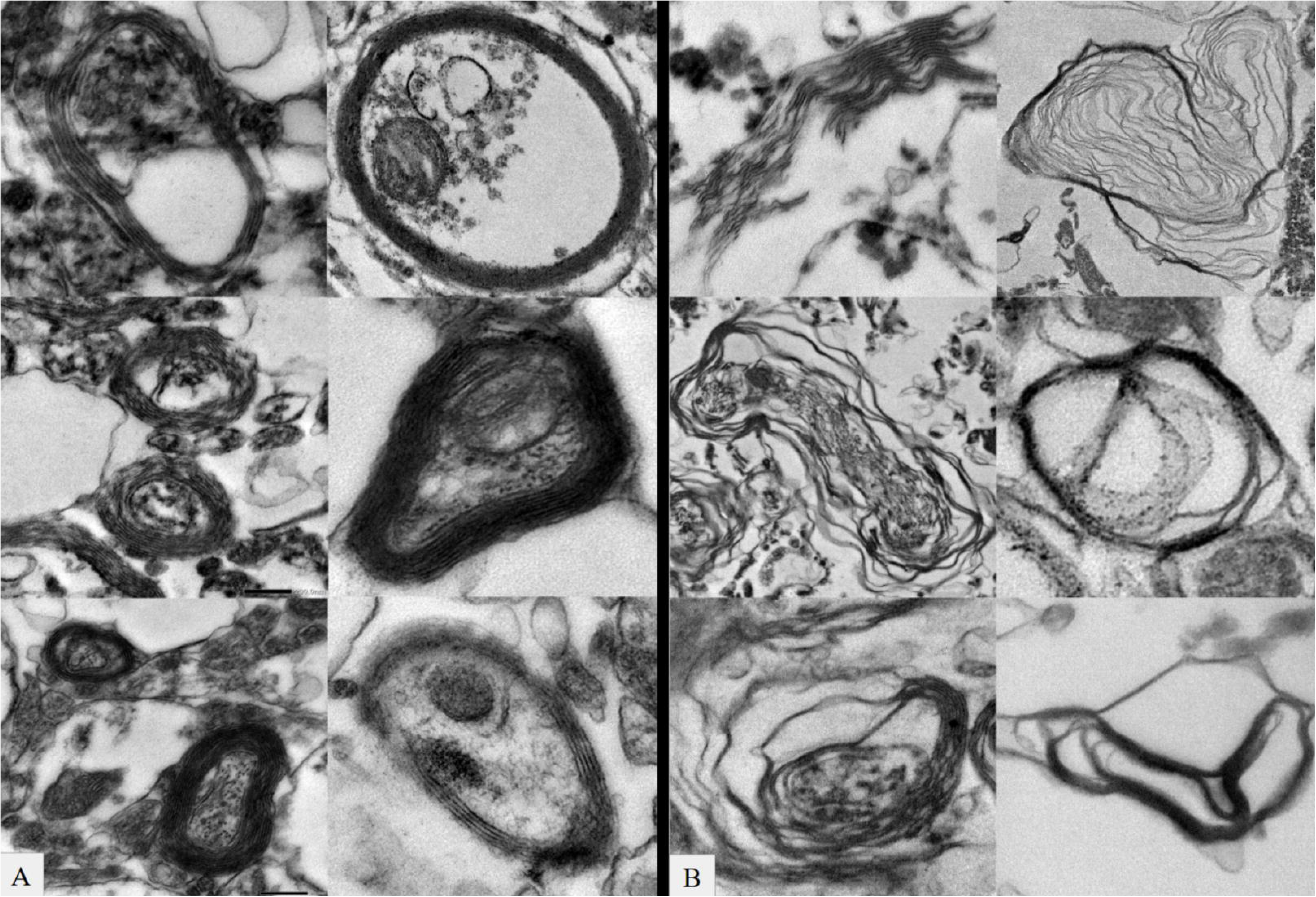
Electron microscopy scans of axons that are either “non-decompacted” (panel A) or “decompacted” (panel B).

### Experimental Design and Statistical Analysis

This secondary data analysis study uses 26 mice including 16 with prolonged exposure to anesthesia and 10 with brief exposure. Information on microbiome group, sex, and exposure to anesthesia can be found in Table 1. Because this was a secondary analysis of existing data, an a priori power analysis was not conducted. A post-hoc sensitivity analysis conducted with G*Power indicates that with an ɑ of 0.05, we had sufficient power (0.80) to dedicate an effect size of d=1.18 (critical t=2.06).

**Table 1.** Sex ratio was similar for mice with prolonged anesthetic exposure and brief anesthetic exposure with a Chi-square value of 0.03, p-value of 0.85, df of 1. Microbiome group did not differ significantly between mice with prolonged anesthetic exposure and brief anesthetic exposure, although there were no SPF animals in the brief anesthetic group; Chi-square value of 3.73, p-value of 0.29, df of 3. HUM1 group was inoculated with a mixed human fecal slurry derived from infants with relatively high levels of *Bifidobacterium*. HUM2 group was inoculated with a mixed human fecal slurry derived from infants with relatively high levels of *Bacteroides*. SPF group received fecal slurry from specific pathogen free mice. GF group received autoclaved fecal slurry from SPF mice and remained in germ-free conditions until behavioral testing.

|  | Prolonged Anesthetic Exposure<br>(N=16) |  | Brief Anesthetic Exposure<br>(N=10) |  |
| --- | --- | --- | --- | --- |
|  | N | % | N | % |
| Sex |  |  |  |  |
| Male | 9 | 60 | 6 | 40 |
| Female | 7 | 63.6 | 4 | 36.4 |
| Group |  |  |  |  |
| SPF | 4 | 100 | 0 | 0 |
| HUM1 | 4 | 44.4 | 5 | 54.6 |
| HUM2 | 4 | 57.1 | 3 | 42.9 |
| GF | 4 | 66.7 | 2 | 33.3 |

Statistical analysis was conducted using GraphPad Prism Version 10.4.0 and R Version 4.5 and Rstudio. The proportion of decompacted axons was calculated for each mouse within each brain region and across all brain regions. Welch’s t-tests were conducted to determine if there were any differences in decompaction between animals with prolonged anesthetic exposure and those without prolonged exposure. These tests were conducted for all axons examined and for axons within each individual brain region. We also conducted a three-way ANOVA to determine if microbiome group or cohort influenced myelin decompaction. A two-way ANOVA was used to determine if sex influenced myelin decompaction. A standard alpha value of 0.05 was used.

## Results

The proportion of decompacted axons was significantly greater in animals exposed to prolonged anesthesia with a p-value of 0.00003642 (df=24) (Figure 3) and an effect size (Cohen’s d) of 1.78.

**Figure 3.**
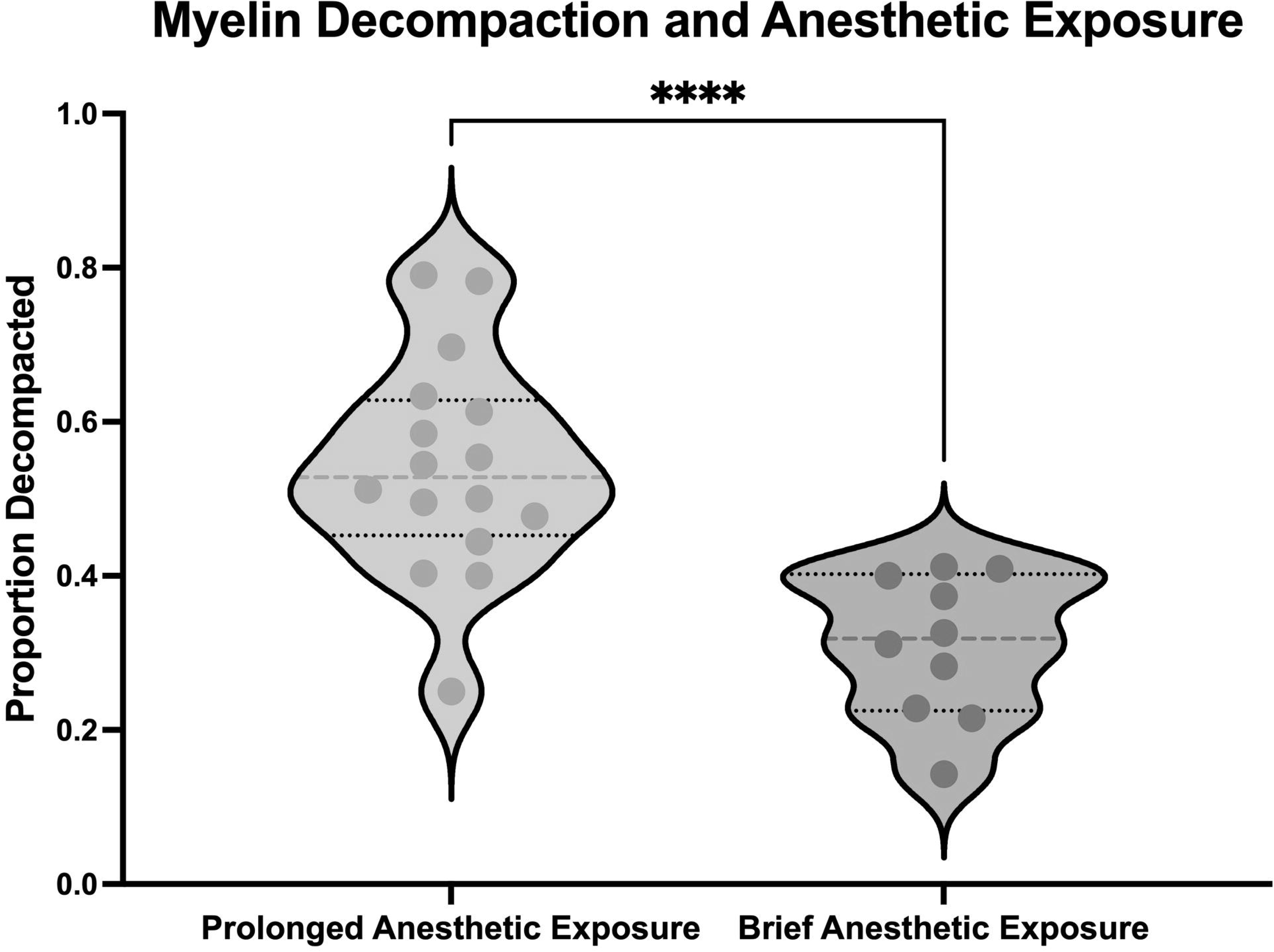
Violin plots showing decompaction for mice with prolonged and brief exposure to anesthetics. *** indicates a p-value of <0.0001

When broken down by region, the difference in decompaction was significant only in the amygdala with a p-value of 0.0011 (df=24) (Figure 4) and an effect size (Cohen’s d) of 2.18.

**Figure 4.**
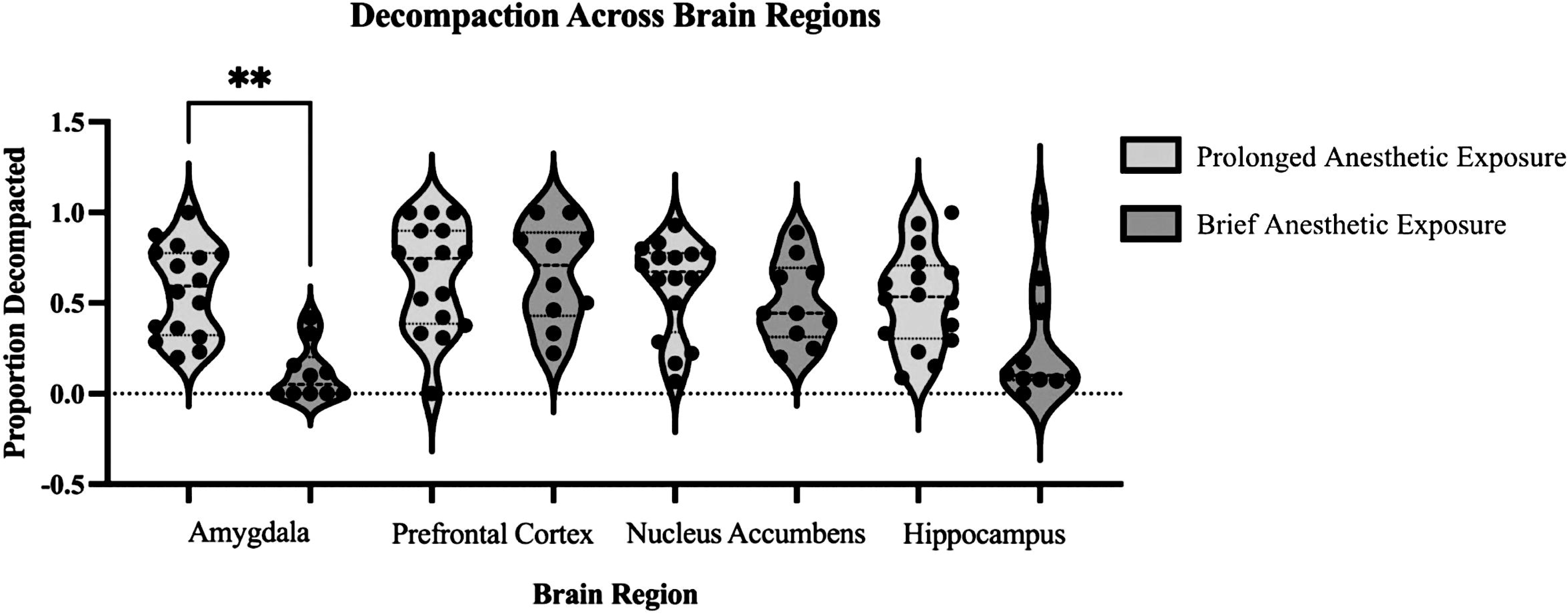
Decompaction across brain regions in study mice. The only region showing significant differences between prolonged anesthetic exposure and brief exposure was the amygdala with a p-value of 0.0011.

Microbiome group, cohort, and sex were not significantly associated with proportion of decompacted axons.

Data supporting these results is available upon request to the corresponding author.

## Discussion

In a secondary examination of a previous study, we found that mice with prolonged exposure to anesthesia during MRI had significantly higher proportions of decompacted axons than mice that did not undergo MRI and were only briefly exposed to anesthesia during humane euthanasia.

This difference seemed to be driven by differences in the amygdala. Here, we discuss potential mechanisms for these results, along with implications and limitations of the study and future directions for work in this field.

### Potential Mechanisms for Myelin Decompaction

Animals in the prolonged exposure group differed from the group without prolonged exposure in three primary ways: (1) they were exposed to isoflurane for an extended period of time (generally 2-3 hours), (2) they were exposed to dexmedetomidine for an extended period of time (generally 2-3 hours), and (3) they were exposed to a high static magnetic field, gradient magnetic fields, and radiofrequency fields used during an MRI scan. We think it is unlikely that the MRI scan itself produced myelin decompaction. Many studies have examined the impact of static magnetic fields on biological tissues without revealing significant effects (Shellock and Crues, 2004). High static magnetic fields may induce acute behavioral changes including circling, suppression of rearing, and the development of conditioned taste aversion, but these appear to be induced by vestibular stimulation and influenced by orientation and restraint within the magnetic field (Houpt et al., 2003; Lockwood et al., 2003; Houpt et al., 2005; Houpt et al., 2007; Houpt et al., 2011). Furthermore, exposure to high static magnetic fields in utero, when the brain is thought to be especially vulnerable to external influences, does not seem to induce long-term changes in neural morphology or behavior(High et al., 2000; Zhu et al., 2011; Hoyer et al., 2012), though one group has reported effects on hippocampal synapses and learning in the Morris water maze (Jiang et al., 2003; Jiang et al., 2004). While gradient magnetic fields can cause peripheral nerve stimulation, we are unaware of any studies documenting axonal damage or behavioral changes in response to gradient stimulation. Radiofrequency fields can produce thermogenic effects within a subject, which could lead to white matter demyelination (Miyamoto et al., 2022). However, we monitored rectal temperatures throughout the MRI scan and did not observe overheating. Consequently, we hypothesize that the myelin decompaction we observed is related to dexmedetomidine exposure and/or isoflurane exposure.

Dexmedetomidine is a highly selective α2 adrenergic receptor agonist that is generally considered to be neuroprotective. For example, dexmedetomidine improves long-term neurological outcomes in neonatal rats with hypoxic-ischemic brain injury by promoting oligodendrocyte genesis and myelination (Xue et al., 2025). However, there is a published report that dexmedetomidine (10 μg) resulted in moderate or severe demyelination in the spinal cord of rabbits when administered via the epidural route (Konakci et al., 2008). The authors hypothesized that this effect was related to vasoconstriction of the medullary spinal vessels and pH of dexmedetomidine. While, neither of these mechanisms seems directly applicable to the current study in which dexmedetomidine was administered subcutaneously via a cannula in the animal’s flank, it is worth noting that dexmedetomidine reduces levels of norepinephrine (Lubinsky and Oster, 2024), a neurotransmitter that increases the intrinsic activity of oligodendrocyte precursors, and that OPC Ca2+ activity is suppressed by in vivo administration of dexmedetomidine (Lu et al., 2023).

Isoflurane is a volatile, halogenated ether that induces anesthesia by inhibiting neurotransmitter-gated ion channels including GABA, glycine, and N-methyl-d-aspartate (NMDA) receptors in the central nervous system (Hawkley et al., 2025). There is a consistent literature documenting adverse effects of isoflurane on the developing brain. Most of this research was conducted in young rodents and non-human primates, where isoflurane induces neural apoptosis, alters levels of synaptic proteins, and, of particular relevance to the current study, disrupts oligodendrocyte development (Liang et al., 2010; Istaphanous et al., 2011; Brambrink et al., 2012; Creeley et al., 2014; Drobish et al., 2016; Kang et al., 2017; Zhao et al., 2020). Studies of isoflurane effects on myelination in adulthood are more limited. One study has reported that repeated exposure to isoflurane in adult male mice results in immediate and lasting motor impairments, along with enduring changes in the microstructural integrity of the corpus callosum (Bajwa et al., 2019).

Another study reported that repeated exposure to sevoflurane, but not single exposure, induced hypomyelination and abnormal ultrastructure of myelin sheath in the prefrontal cortex (PFC) of adult male mice and exacerbated demyelination in the cuprizone-induced model of multiple sclerosis (MS) (Zhang et al., 2025). Sevoflurane, like isoflurane, is a halogenated ether used for inhalation anesthesia. On the other hand, isoflurane preconditioning has been reported to protect against inflammation, cerebral lipid peroxidation, and histologic injury in a rat model of focal cerebral ischemia (Bedirli et al., 2012).

One key mechanism by which isoflurane could impact myelin decompaction is neuroinflammation. Many neurological diseases are characterized by neuroinflammation, which induces blockade of myelin formation, loss of mature myelin, and/or suppression of remyelination via effects on oligodendrocytes (Festa et al., 2025). Isoflurane can activate the inflammatory mTOR pathway, and previous studies have linked this activation to oligodendrocyte disruption in the hippocampal fibria of infant mice (Kang et al., 2017).

Inflammation can also damage the myelin sheath directly via macrophage invasion into and below myelin sheaths allowing macrophage-derived proteinases to degrade myelin proteins (Wisniewski and Bloom, 1975). Another potential mechanism is via Wnt/β-catenin signaling pathways (Ma et al., 2017). Isoflurane influences expression levels of Wnt3a, GSK 3β and β- catenin, and excessive Wnt activity disrupts developmental and regenerative myelination (Norton et al., 1978). A third potential mechanism is via the p75NTR pathway. While this pathway has primarily been investigated as a mediator of isoflurane-induced apoptosis (Lemkuil et al., 2011; Schilling et al., 2017; Sohn et al., 2017), it has recently been linked to oligodendrocyte production and maturation during myelin formation (Joshi et al., 2023).

### Region-Specific Environmental Sensitivity

When broken down by region, the difference in decompaction was significant only in the amygdala and not in the prefrontal cortex, nucleus accumbens, or hippocampus. This was somewhat surprising. While detrimental effects of isoflurane on the amygdala have been reported, there are also reports of detrimental effects on the hippocampus, nucleus accumbens, and in cortex (Liang et al., 2010; Brambrink et al., 2012; Creeley et al., 2014; Drobish et al., 2016; Long Ii et al., 2016; Ma et al., 2017; Li et al., 2024). Perhaps there is less regional specificity in the developing brain, due to widespread and rapid oligodendrogenesis during this period, though it must be noted that myelination is a protracted and plastic process continuing into adulthood (Sturrock, 1980; El Waly et al., 2014; de Faria et al., 2021). It is interesting to note that there are distinct GABAergic neurons activated by isoflurane in the amygdala that have been shown to be responsible for the analgesic effect of the anesthetic (Hua et al., 2020). Perhaps this also contributes to the unique vulnerability of the amygdala in our study.

### Study Limitations

Because this study was a secondary analysis of existing data, there are several limitations. First, the study design did not allow us to distinguish whether the effects we observed were due to isoflurane, dexmedetomidine, or MRI exposure. Second, because animals were sacrificed shortly after exposure, we do not know if this protocol induced persistent changes in myelination or altered animals’ cognition and behavior. In the future, several groups of mice should be kept alive at varying time points post-exposure to test for behavioral and cognitive changes and to determine if decompaction resolves over time. Lastly, no analyses were conducted to elucidate potential mechanisms for decompaction; future studies could incorporate measures of inflammation, Wnt/β-catenin signaling, and the p75NTR pathway.

### Study Implications and Future Directions

This is one of the first studies demonstrating adverse effects of prolonged exposure to isoflurane and dexmedetomidine on myelination in adult mice. This has significant implications for medicine and scientific research, as these anesthetics are commonly used in both. However, further studies must be done to replicate these findings, as well as to explore the long-term impacts of prolonged anesthetic exposure on behavior and myelination. This specific anesthetic protocol was selected for the parent study because it was considered especially appropriate for acquiring resting-state fMRI data in mice. Functional magnetic resonance imaging (fMRI) is commonly used in rat and mice models to study mechanisms underlying physiological as well as pathological brain functions. Researchers have drawn attention to the diversity of currently used anesthetic protocols and urged the field to pursue standardization of doses, routes and timing of administration (Steiner et al., 2021). As the field moves toward standardization, it will be important to consider not only the impact of protocol on BOLD fMRI outcomes, but also potential effects on brain structure. Overall, while MRI holds great promise for conducting studies of the brain in rodents, researchers need to be cognizant that anesthetic protocols could impact results and must be selected with awareness. Regarding implications for veterinary medicine, our results add to a growing body of literature suggesting enhanced caution regarding use of sedatives for procedures that could be conducted on conscious animals such as routine collection of blood samples or annual physical examinations. Regarding human medicine, if the effects we observed translate to human beings, this would have important implications for clinical practice and for control of occupational exposures.

## Conflict of Interest

The authors declare no competing financial interests

### Acknowledgement statement

We thank the staff at MSU’s Center for Advanced Microscopy, especially Alicia Withrow, for generating the electron microscopy images and Erik Shapiro, Faculty Lead for MSU’s Advanced Molecular Imaging Facility in the Institute for Quantitative Health Science and Engineering, where the MRIs were acquired. Funding was provided by MSU through start-up funds to Dr. Knickmeyer

